# Quantifying protein oligomerization in living cells: A systematic comparison of fluorescent proteins

**DOI:** 10.1101/311175

**Authors:** Valentin Dunsing, Madlen Luckner, Boris Zühlke, Roberto Petazzi, Andreas Herrmann, Salvatore Chiantia

**Affiliations:** Institute for Biochemistry and Biology, Universität Potsdam, Karl-Liebknecht-Str. 24-25, 14476 Potsdam; Institute for Biology, IRI Life Sciences, Humboldt-Universität zu Berlin, Invalidenstraße 42, 10115 Berlin

**Author notes:** co-first authors (Valentin Dunsing, Madlen Luckner).

## Abstract

Fluorescence fluctuation spectroscopy has become a popular toolbox for non-disruptive studies of molecular interactions and dynamics in living cells. The quantification of e.g. protein oligomerization and absolute concentrations in the native cellular environment is highly relevant for a detailed understanding of complex signaling pathways and biochemical reaction networks. A parameter of particular relevance in this context is the molecular brightness, which serves as a direct measure of oligomerization and can be easily extracted from temporal or spatial fluorescence fluctuations. However, fluorescent proteins (FPs) typically used in such studies suffer from complex photophysical transitions and limited maturation, potentially inducing non-fluorescent states, which strongly affect molecular brightness measurements. Although these processes have been occasionally reported, a comprehensive study addressing this issue is missing.

Here, we investigate the suitability of commonly used FPs (i.e. mEGFP, mEYFP and mCherry), as well as novel red FPs (i.e. mCherry2, mRuby3, mCardinal, mScarlet and mScarlet-I) for the quantification of oligomerization based on the molecular brightness, as obtained by Fluorescence Correlation Spectroscopy (FCS) and Number&Brightness (N&B) measurements in living cells. For all FPs, we measured a lower than expected brightness of FP homo-dimers, allowing us to estimate, for each fluorescent label, the probability of fluorescence emission in a simple two-state model. By analyzing higher FP homo-oligomers and the Influenza A virus Hemagglutinin (HA) protein, we show that the oligomeric state of protein complexes can only be accurately quantified if this probability is taken into account. Further, we provide strong evidence that mCherry2, an mCherry variant, possesses a superior apparent fluorescence probability, presumably due to its fast maturation. We finally conclude that this property leads to an improved quantification in fluorescence cross-correlation spectroscopy measurements and propose to use mEGFP and mCherry2 as the novel standard pair for studying biomolecular hetero-interactions.

## INTRODUCTION

A large variety of biological processes rely on transport and interactions of biomolecules in living cells. For a detailed understanding of these events, minimally invasive techniques are needed, allowing the direct quantification of inter-molecular interactions in the native cellular environment. In recent years, fluorescence fluctuation spectroscopy (FFS) approaches have been often used to fulfill this task^1–6^. FFS is based on the statistical analysis of signal fluctuations emitted by fluorescently labeled molecules. While the temporal evolution of such fluctuations provides information about dynamics, the magnitude of the fluctuations contains information about molecule concentration and interactions (i.e. oligomeric state). In order to probe the oligomerization of a protein of interest directly in living cells, the molecular brightness (i.e. fluorescence count rate per molecule) of a genetically fused fluorescent protein (FP) is analyzed^4,5,7^. Comparison to a monomeric reference allows the quantification of the number of FPs within a protein complex, i.e. its oligomeric state. This analysis can be performed with different experimental methods, e.g. Fluorescence Correlation Spectroscopy (FCS)^1,8^, Photon Counting Histogram (PCH)^4,9^, Number&Brightness analysis (N&B)^2,7^ or subunit counting^10,11^.

Measuring the oligomeric state from the number of fluorescent labels, it is often assumed that all FPs emit a fluorescence signal. However, various *in vitro* studies of FPs revealed complex photophysical properties such as: long-lived dark states of green FPs^12–15^, transitions between different brightness states (e.g. YFP^16^, mCherry^17^) and flickering^18^. Additionally, limited maturation was reported for FPs expressed in cells^19^. All together, these observations challenge the suitability of FPs for quantitative brightness analysis^5^. In this context, partially contradicting results are reported: studies performing subunit counting typically indicate apparent fluorescence probability (pf) values of 50-80%^10,11,20,21^ for GFPs. Very few investigations utilizing FFS approaches report similar values^22,23^, while very often it is simply assumed that all FPs are fluorescent. For commonly used red FPs (mainly RFP and mCherry), published results tend to agree, consistently reporting low p_f_ values (ca. 20-40%)^24,25^, with only few exceptions^17^.

Notably, many investigations would profit from systematic controls testing the presence of non-fluorescent labels, but so far only few studies take explicitly into account the role of the p_f_ in the exact quantification of protein-protein interaction^5,11,22,26^. Importantly, oligomerization data are prone to severe misinterpretations if non-fluorescent labels are not taken into consideration. To our knowledge, this is the first report systematically comparing non-fluorescent states and associated p_f_ for various FPs in one-photon excitation. We found significant amounts of non-fluorescent FPs in different cell types and compartments, and we determined the p_f_ for each FP. With appropriate corrections, we were able to correctly determine the oligomeric state of the homo-trimeric Influenza A virus Hemagglutinin glycoprotein, for the first time directly in living cells, as a proof of principle.

To investigate multiple interacting molecular species simultaneously, multicolor FFS analysis is often performed. For example, protein hetero-interactions can be quantified via fluorescence cross-correlation approaches^1,27^, even in living multicellular organisms^6,28^. Such methods require well-performing FPs with spectral properties distinguished from the typically used mEGFP. Therefore, current FP development focuses on red and far-red FPs^29^. Nevertheless, the p_f_ for these proteins, although playing a fundamental role in brightness and cross-correlation analysis, has not been systematically investigated yet. We therefore screened different red FPs for the presence of non-fluorescent states, and found that mCherry2, a new mCherry variant, possesses superior properties compared to all other tested red FPs, i.e. mCherry, mCardinal, mRuby3, mScarlet and mScarlet-I. Additionally, by performing FCCS measurements of FP hetero-dimers, we show that mCherry2 improves the quantification of the spectral crosscorrelation compared to mCherry and propose to use mEGFP and mCherry2 as a novel standard FP pair for hetero-interaction studies.

## Results

### The brightness of homo-dimers of conventional FPs is lower than double the brightness of monomers

In an ideal case, i.e. if all fluorophores within an oligomer were fluorescent, a homo-dimer would emit twice as many photons as a monomer. We expressed several FPs in the cytoplasm of HEK 293T cells and performed FFS measurements. We found that the brightness of homo-dimers (normalized to the brightness of the corresponding monomer) for three widely used FPs, namely mEGFP (ε_dimer_=1.69 ± 0.05), mEYFP (ε_dimer_ =1.63 ± 0.05) and mCherry (ε_dimer_ =1.41 ± 0.04), are generally lower than two, indicating the presence of non-fluorescent proteins. The effect is particularly pronounced for mCherry (Figure 1A) and does not depend on the specific FFS method used or cellular localization, as shown by comparing the results from N&B, pFCS (in cytoplasm and nucleus) and sFCS (for FPs associated to the plasma membrane (PM)) (Figure 1A, B). Interestingly, we observed a 10% lower brightness for FP monomers within the nucleus compared to the cytoplasmic fraction (Figure S1A). Furthermore, we measured homo-dimer brightness values of mEGFP and mCherry in different cell lines (HEK 293T, A549, CHO, HeLa) and obtained comparable values in all cell types for the same FP (Figure S1B, C).

**Figure 1.**
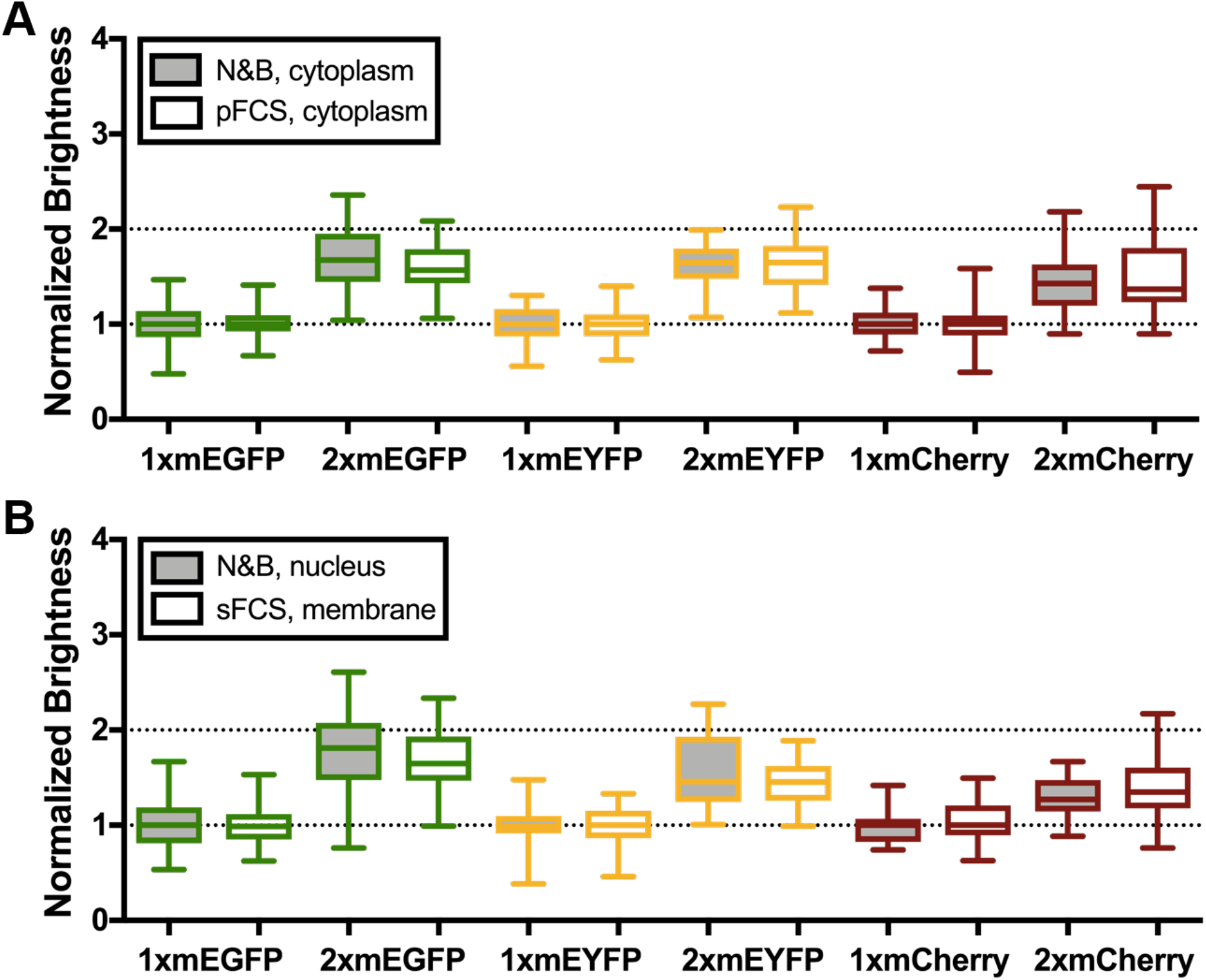
Brightness comparison of different FPs in living HEK 293T cells. **A**: Box plots of normalized molecular brightness of mEGFP, mEYFP and mCherry monomers and homo-dimers in HEK 293T cells, measured via N&B (grey) or pFCS (white). Monomer and dimer constructs are labeled as “1x” and “2x”, respectively. Data were pooled from at least three independent experiments (N&B/pFCS: 1xmEGFP: n=47/39 cells, 2xmEGFP: n=48/38 cells, 1xmEYFP: n=33/37 cells, 2xmEYFP: n=32/39 cells, 1xmCherry: n=50/35 cells, 2xmCherry: n=53/34 cells). B: Normalized molecular brightness of mEGFP, mEYFP and mCherry monomers and homo-dimers in the nucleus (grey) and plasma membrane (PM, white) of HEK 293T cells, measured with N&B (nucleus) and scanning FCS (PM). For PM measurements, myristoylated-palmitoylated 1xmEGFP (mp 1xmEGFP), mp 2xmEGFP, mp 1xmEYFP, mp 2xmEYFP, GPI 1xmCherry and GPI 2xmCherry constructs were expressed. See Methods section for a description of the investigated FP constructs. Data were pooled from at least three independent experiments (nucleus: 1xmEGFP: n=47 cells, 2xmEGFP: n=48 cells, 1xmEYFP: n=30 cells, 2xmEYFP: n=32 cells, 1xmCherry: n=32 cells, 2xmCherry: n=37 cells; PM: mp 1xmEGFP: n=55 cells, mp 2xmEGFP: n=55 cells, mp 1xmEYFP: n=28 cells, mp 2x mEYFP: n=28 cells, GPI 1xmCherry: n=38 cells, GPI 2xmCherry: n=38 cells).

The maturation time of FPs might influence the fraction of non-fluorescent proteins and this, in turn, may be dependent on the temperature at which experiments are performed^30^. For this reason, we compared the homo-dimer brightness of mEGFP at 23°C and 37°C, but observed negligible differences (Figure S1D).

Taken together, our results demonstrate that the effect of non-fluorescent states on brightness quantification for mEGFP, mEYFP and mCherry is mainly a fluorophore-inherent property and is not strongly influenced by the tested experimental conditions.

### The oligomeric state of mEGFP homo-oligomers is correctly determined by using a simple correction scheme for non-fluorescent states

Based on the observed non-fluorescent protein fractions for mEGFP, mEYFP and mCherry, we investigated whether it is possible to nevertheless correctly determine the oligomeric state of higher-order oligomers. To this aim, we expressed mEGFP-homo-oligomers of different sizes: 1xmEGFP, 2xmEGFP, 3xmEGFP and 4xmEGFP (i.e. monomers to tetramers). We then performed pFCS measurements in the cytoplasm of living A549 cells (Figure 2A).

**Figure 2.**
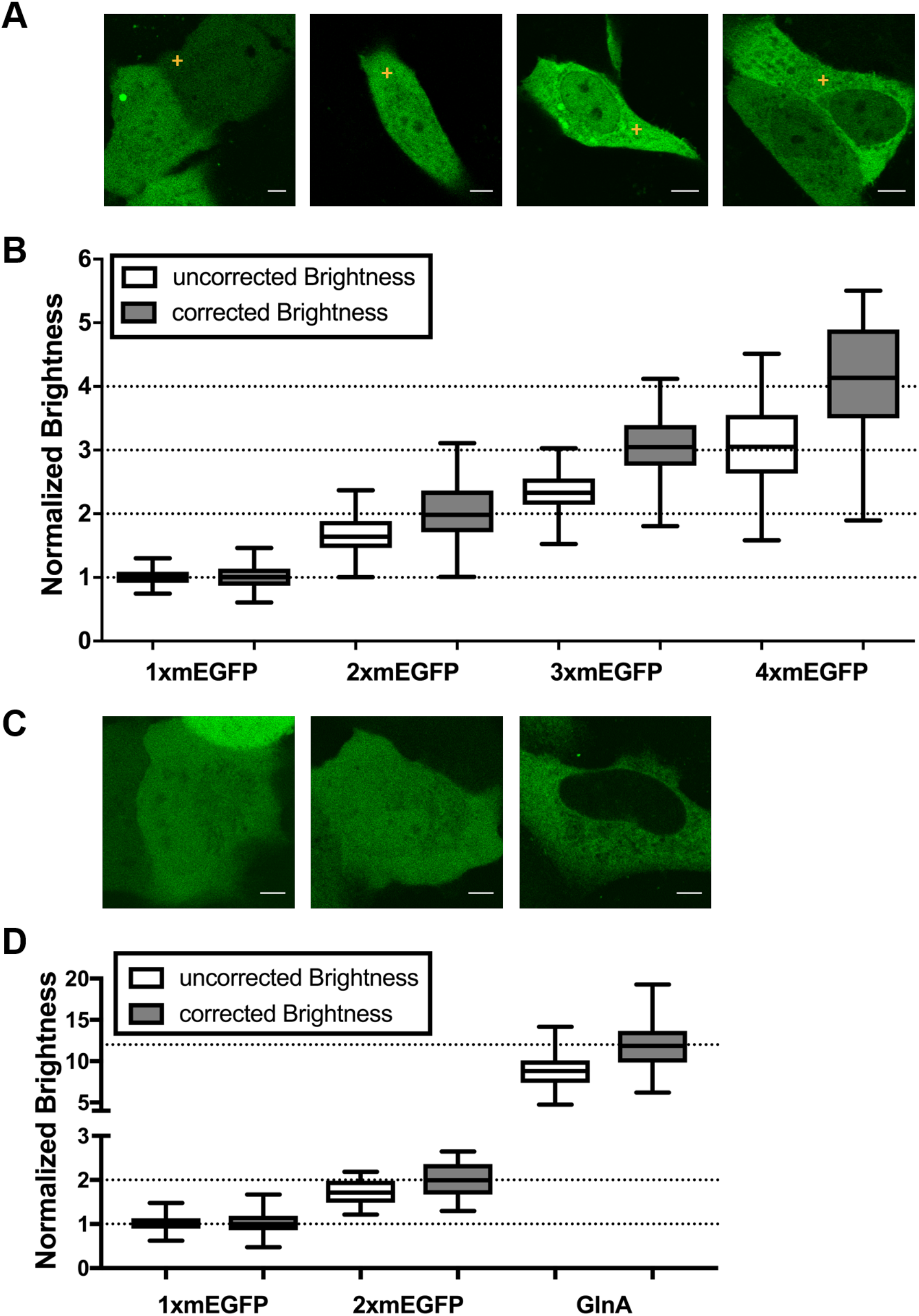
Brightness analysis of mEGFP homo-oligomers. **A**: Representative images of A549 cells expressing 1xmEGFP, 2xmEGFP, 3xmEGFP and 4xmEGFP, from left to right. Yellow crosses indicate the positions of the pFCS scan point. Scale bars are 5 μm. **B**: Box plots of normalized molecular brightness obtained from pFCS analysis, pooled from at least three independent experiments (1xmEGFP: n=52 cells, 2xmEGFP: n=42 cells, 3xmEGFP: n=43 cells, 4xmEGFP: n=59 cells) before correction (white) and after correction (grey). First, a normalization of the uncorrected brightness data was performed using the brightness value of 1xmEGFP. Second, a correction was performed as described in the Methods section, using a p_f_ of 0.65, as obtained from measurements on 2xmEGFP. **C**: Representative images of U2OS cells expressing 1xmEGFP, 2xmEGFP and GlnA-mEGFP (GlnA). Scale bars are 5 μm. **D**: Box plots of normalized molecular brightness obtained from N&B analysis, pooled from three independent experiments (1xmEGFP: n=34 cells, 2xmEGFP: n=35 cells, GlnA: n=41 cells) before correction (white) and after correction (grey). After normalization using the brightness value of 1xmEGFP, a correction was performed using a p_f_ of 0.72, as obtained from measurements on 2xmEGFP.

We observed brightness values consistently lower than those expected. For example, the obtained tetramer brightness (ε_tetramer_=3.01 ± 0.08) is closer to the theoretical trimer brightness value (Figure 2B, white boxes). Hence, we performed a brightness correction based on a simple two-state model^11,31^, taking into account the probability that each FP subunit emits a fluorescence signal. The p_f_ values were determined from the brightness of 2xmEGFP (ε_dimer_=1.65 ± 0.06, p_f_=0.65). Thus, we were able to correctly determine the oligomeric state of all mEGFP-homo-oligomers investigated in this study (Figure 2B, grey boxes). Consistent with the brightness data, pFCS analysis revealed an increase of the diffusion times with increasing homo-oligomer size (SI and Figure S2A-C).

Furthermore, to extend our investigation to larger protein complexes, we performed N&B measurements on U2OS cells expressing the dodecameric *E.coli* glutamine synthetase (GlnA)^32^. We measured an average normalized brightness of ε_12-mer_=8.8 ± 0.3. However, after correction for non-fluorescent mEGFP subunits (ε_dimer_=1.72 ± 0.05, p_f_=0.72), we obtained an oligomeric state of ε_12-mer_=11.9 ± 0.4, confirming the expected 12-mer structure of the GlnA complex.

Overall, these results highlight the importance of performing control experiments with suitable homo-oligomers for brightness-based oligomerization studies and demonstrate that the simple correction for non-fluorescent states presented here produces reliable results.

### Influenza A virus hemagglutinin forms homo-trimers in the plasma membrane

We next verified whether the above-mentioned simple two-state brightness correction provides reliable quantitative results in a biologically relevant context. We analyzed an mEGFP-fused version of the Influenza A virus hemagglutinin (HA-wt-mEGFP), a biochemically well-characterized trimeric transmembrane protein^33,34^. To this aim, we expressed the fluorescent construct in living HEK 293T cells and performed sFCS measurements (Figs. 3A and S3) across the PM. After correction for the non-fluorescent FPs contribution, we obtained an average normalized brightness of ε_HA_=3.17±0.12 (Fig 3B), in line with the expected trimeric structure of HA-wt-mEGFP. We further investigated an HA-TMD mutant, in which the HA ectodomain is replaced by mEGFP on the extracellular side. This construct was shown to localize as HA-wt in the PM, but in a dimeric form^35^. The observed brightness of HA-TMD-mEGFP was significantly lower than that of HA-wt-mEGFP (Figure 3B). After correcting for non-fluorescent FPs, we found an average normalized brightness of ε_HA-TMD_=1.82 ± 0.07, compatible with the presence of a large dimer fraction.

**Figure 3.**
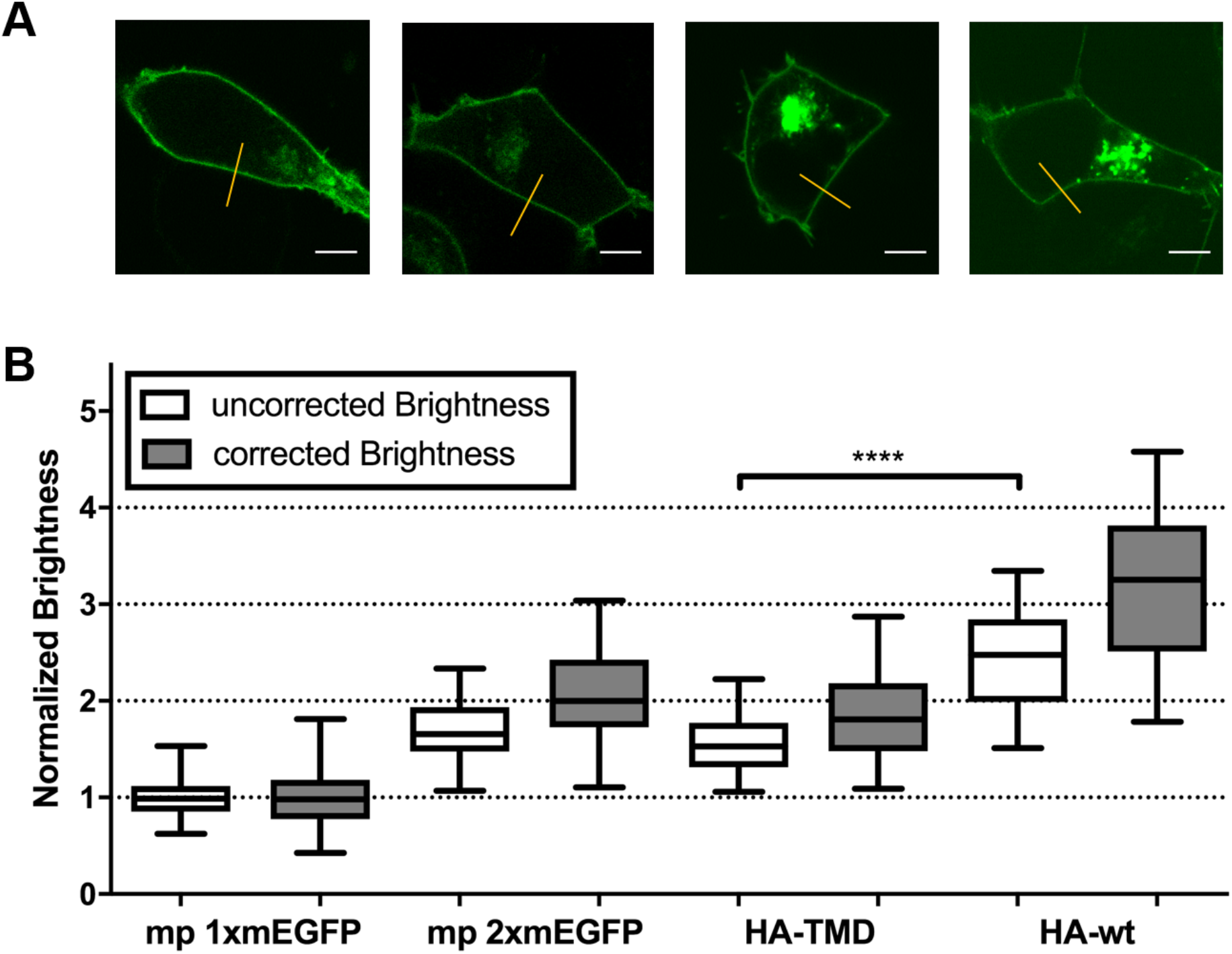
Oligomerization of Influenza A virus hemagglutinin (HA) protein measured with sFCS. **A**: Representative images of HEK 293T cells expressing mp 1xmEGFP, mp 2xmEGFP, HA-TMD-mEGFP (HA-TMD) and HA-wt-mEGFP (HA-wt), from left to right. Yellow lines indicate sFCS scan lines. Scale bars are 5 μm. **B**: Box plots of normalized molecular brightness obtained from sFCS analysis, pooled from at least three independent experiments (mp 1xmEGFP: n=55 cells, mp 2xmEGFP: n=54 cells, HA-TMD: n=37 cells, HA-wt: n=36 cells) before correction (white) and after correction (grey) with p_f_=0.65 obtained from mp 2xmEGFP measurements. **** indicates significance with p<0.0001, obtained by using a two-tailed Mann-Whitney test.

In summary, these results clearly demonstrate that a simple two-state model for FFS-derived brightness data correction allows precise quantification of the oligomeric state of proteins in living cells.

### The mCherry variant “mCherry2” has a superior performance in FFS measurements, compared to other red fluorescent proteins

In order to extend brightness measurements to the investigation of hetero-interactions, FPs with spectral properties different from those of mEGFP are needed. Typically, red FPs are well suited for this task since spectral overlap with mEGFP is low, reducing the possibility of FRET or cross-talk. However, for the typically used mCherry, we and others^25^ observed a high fraction of non-fluorescent states, i.e. only ca. 40% of the proteins were fluorescent. In order to identify red FPs with higher p_f_, we screened the recently developed FPs mCherry2^36^, mCardinal^37^, mRuby3^38^, mScarlet^39^ and mScarlet-I^39^. We performed bleaching and N&B measurements of monomers and homo-dimers, expressed in HEK 293T cells. Notably, we observed strong photobleaching for mRuby3, mScarlet and mScarlet-I (Figure 4A, Table 1) compared to the other three tested FPs. Therefore, N&B measurements on these proteins were conducted at lower excitation powers. This reduces their effective brightness, e.g. only 1 kHz for mRuby3, compared to the theoretically three-fold higher brightness when interpolated to the same laser powers used for mCherry, mCherry2 and mCardinal (Figure 4B). All other FPs exhibit minor difference in the effective brightness ranging from 1.5 kHz (mCherry2) to 2.2 kHz (mCardinal, mScarlet) in our experimental conditions. However, when comparing the normalized homo-dimer brightness, we found strong differences between mCherry2 and the other FPs. We estimated a p_f_ of 0.71 for mCherry2, which is ~1.8-fold higher than that of mCherry (p_f_=0.41) and mScarlet (p_f_=0.40), while mCardinal and mRuby3 show very low p_f_ values of only 0.24 and 0.22, respectively. Notably, mScarlet-I also features a high p_f_(0.63), but still suffers from considerable photobleaching, even at lower excitation powers (Figure 4C, Table 1).

**Figure 4.**
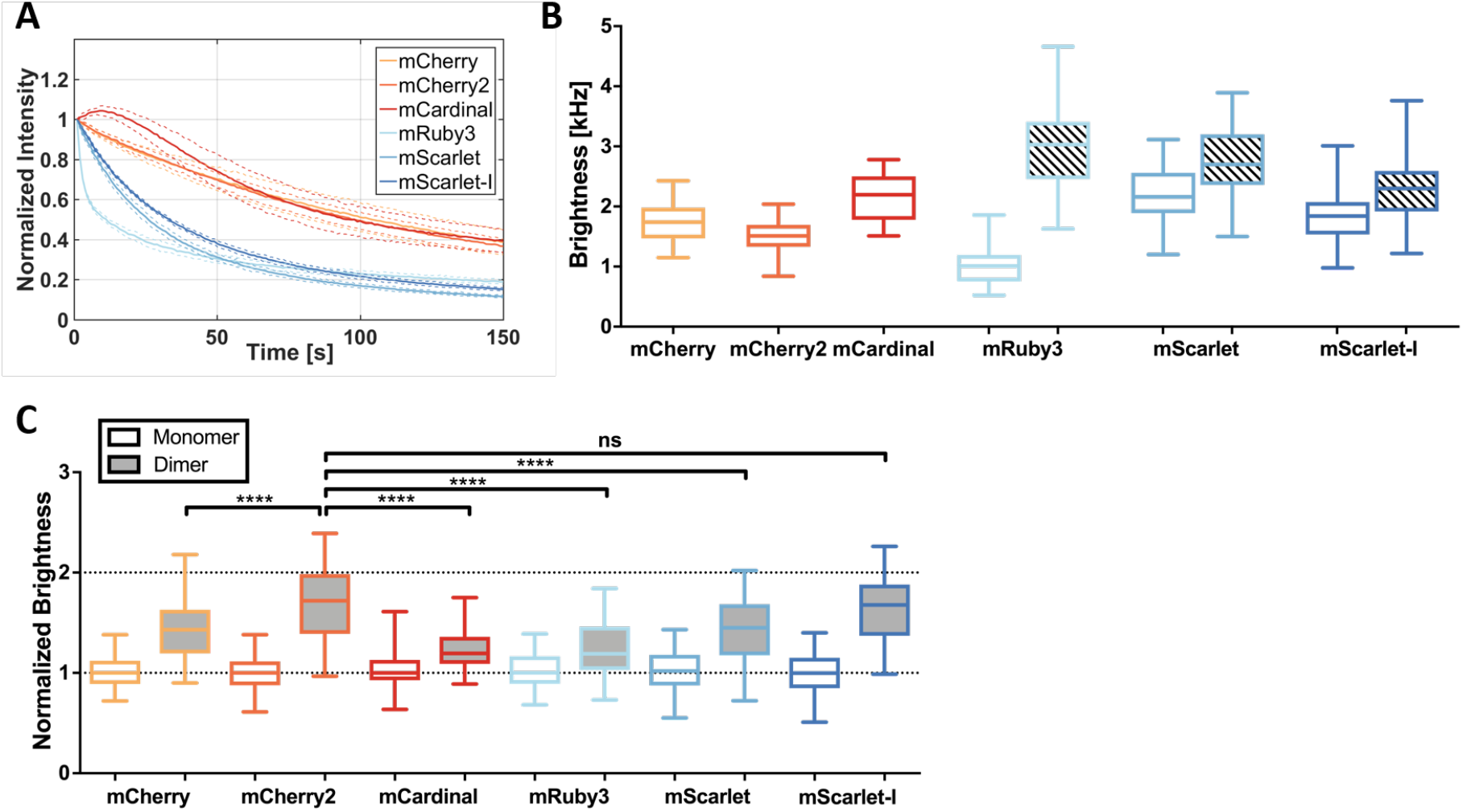
Comparison of different red fluorescent proteins (FPs) in bleaching and number and brightness (N&B) measurements. **A**: Bleaching curves of different red FPs, mCherry, mCherry2, mCardinal, mRuby3, mScarlet and mScarlet-I, expressed in HEK 293T cells, obtained in three independent N&B measurements of 18 cells each, with 19.6μW laser power (four-fold compared to standard N&B settings). Solid lines show average curves, dashed lines show mean±std. **B**: Box plots of molecular brightness of different red FP monomers expressed in HEK 293T cells, measured with N&B at 4.9μW (mCherry, mCherry2, mCardinal), 3.9μW (mScarlet, mScarlet-I) or 1.6μW (mRuby3) laser power in three independent experiments (mCherry: n=51 cells, mCherry2: n=49 cells, mCardinal: n=32 cells, mRuby3: n=33 cells, mScarlet: n=36 cells, mScarlet-I: n=34 cells) (white boxes). The different excitation laser powers were required to avoid strong bleaching for the less photostable FPs (e.g. mRuby3). The shaded boxes for mRuby3, mScarlet and mScarlet-I show brightness values interpolated to 4.9μW laser power, assuming a linear increase of the brightness with the excitation laser power. **C**: Box plots of normalized molecular brightness of red FP monomers (white boxes) and dimers (grey boxes). Data represent results of three independent experiments (1xmCherry: n=50 cells, 2xmCherry: n=53 cells, 1xmCherry2: n=49 cells, 2xmCherry2: n=54 cells, 1xmCardinal: n=42 cells, 2xmCardinal: n=42 cells, 1xmRuby3: n=33 cells, 2xmRuby3: n=31 cells, 1xmScarlet: n=36 cells, 2xmScarlet: n=41 cells, 1xmScarlet-I: n=34 cells, 2xmScarlet-I: n=39 cells). **** indicates statistical significance compared to mCherry2 with p<0.0001, ns indicates no statistical significance, obtained by using a two-tailed Mann-Whitney test.

**Table 1:** Characteristics of all investigated red FPs.

| FP | $t_{1/2}^*$ [s] | $\epsilon_{N\&B}$ [kHz] | Bleaching<sub>N&B</sub> [%] | $\epsilon^{**}$ [kHz] | $p_f$ |
| --- | --- | --- | --- | --- | --- |
| mCherry | 104 | 1.7 | 6.2 | 1.7 | 0.41 |
| mCherry2 | 99 | 1.5 | 4.3 | 1.5 | 0.71 |
| mCardinal | 97 | 2.2 | -3.5 | 2.2 | 0.24 |
| mRuby3 | 12 | 1.1 | 37.5 | 3.1 | 0.21 |
| mScarlet | 29 | 2.2 | 31.7 | 2.7 | 0.40 |
| mScarlet-I | 34 | 1.9 | 29.0 | 2.4 | 0.63 |

\* Measured at four times higher laser power (19.6 μW) than used in N&B measurements
\*\* Average molecular brightness in N&B, interpolated to the same laser power (4.9 μW) for all red FPs

The superior performance of mCherry2 was confirmed in other cell types, as we consistently observed a reproducible difference from mCherry (Figure S4 A). Moreover, we compared the homo-dimer brightness of mCherry2 at 23°C and 37°C and, similarly to mEGFP, observed only negligible variations (Figure S4 B).

We therefore conclude that mCherry2 exhibits cell type- and temperature-independent, superior properties in the context of FFS measurements, compared to all the other tested red FPs.

### Quantification of hetero-interactions via fluorescence cross-correlation spectroscopy is improved by using mCherry2

Cross-correlation techniques (e.g. FCCS^1^, ccN&B^2^, RICCS^3^) are powerful methods for the investigation of protein hetero-interactions. These techniques are based on the analysis of simultaneous fluorescence fluctuations emitted by co-diffusing, spectrally distinct labeled molecules. In protein-protein interaction studies, fusion proteins with mEGFP and mCherry or RFP are typically used for this purpose^1,2,6,40^. However, due to the presence of non-fluorescent states, only a fraction of protein hetero-complexes simultaneously emits fluorescence in both channels (i.e. many complexes will contain fluorescent green proteins and non-fluorescent red proteins). This factor has to be taken into account when calculating e.g. dissociation constants from cross-correlation data^25^. Given the superior p_f_ of mCherry2 compared to other red FPs, we hypothesized that mCherry2 would improve the quantification of cross-correlation data, since more complete fluorescent protein-complexes should be present. To test this hypothesis, we performed pFCCS experiments with mCherry-mEGFP and mCherry2-mEGFP hetero-dimers in the cytoplasm of living A549 cells. As presumed, we observed a higher auto-correlation function (ACF) amplitude G in the red than in the green channel (G_g_/G_r_=0.65±0.03, Figure 5A,C) for mCherry-mEGFP, indicating that the apparent concentration of mCherry is ca. 1.5-fold lower than that of mEGFP (i.e. in a significant fraction of hetero-dimers, only mEGFP is fluorescent). This is in agreement with the expected relative amount of hetero-dimers containing fluorescent mEGFP and/or mCherry, based on the above-mentioned p_f_ values. Furthermore, we expect ~27% of hetero-dimers to carry both fluorescent mEGFP and mCherry (see SI related to Figure S5).

**Figure 5.**
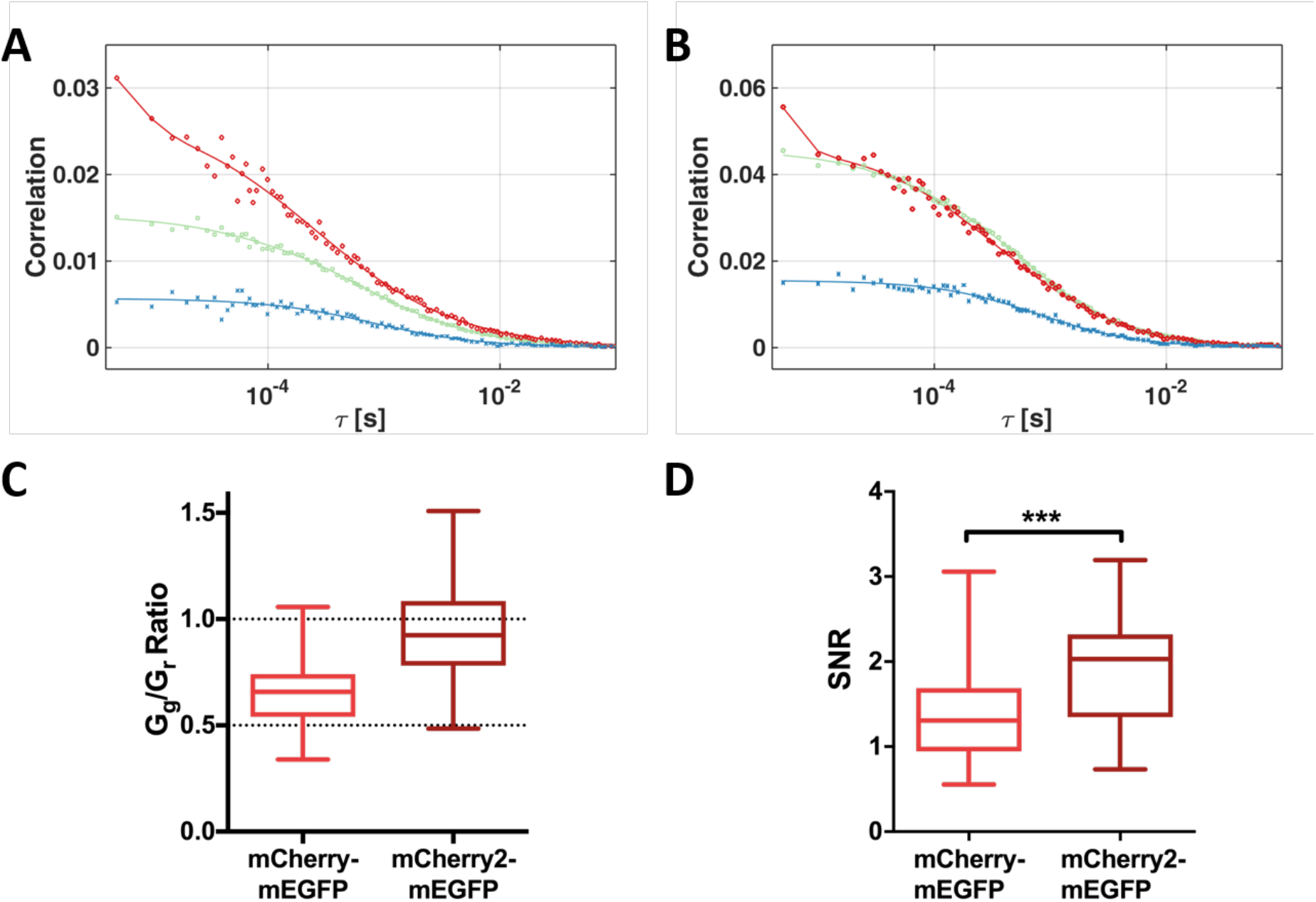
Cross-correlation measurements of mCherry-/mCherry2-mEGFP hetero-dimers. **A** and **B**: Representative correlation functions and fit curves for pFCCS measurements of mCherry-mEGFP (A) and mCherry2-mEGFP (B) heterodimers in A549 cells. Green, ACF in green channel (mEGFP); red, ACF in red channel (mCherry (A), mCherry2 (B)); blue, CCF calculated for both spectral channels. Fit curves (solid lines) were obtained from fitting a three-dimensional anomalous diffusion model to the data. **C**: Box plots of amplitude ratios of the green to red ACFs for mCherry-mEGFP (n=35 cells) and mCherry2-mEGFP (n=32 cells) pooled from three independent experiments performed in A549 cells. **D**: Box plots of signal to noise ratios (SNR) of the CCFs for mCherry-mEGFP and mCherry2-mEGFP hetero-dimers, calculated from pFCCS measurements in A549 cells, described in (C).

For mCherry2-mEGFP in contrast, the amplitudes of the ACFs in the red and green channel were comparable (G_g_/G_r_=0.97±0.05, Figure 5B,C), indicating, as expected, similar apparent concentrations of mCherry2 and mEGFP (see SI related to Figure S5). Also, the relative amount of hetero-dimers carrying fluorescent mEGFP and mCherry2 is estimated to be ~42%, i.e. 1.5-fold more fully-fluorescent complexes than for mCherry-mEGFP. On the other hand, the expected cross-correlation values for mCherry- and mCherry2-mEGFP should be similar, which is confirmed by our data (Figure S5). Nevertheless, the 1.5-fold higher relative fraction of fully-fluorescent hetero-dimers with mCherry2 should improve the quality of crosscorrelation data. We therefore compared the signal-to-noise (SNR) ratio of the measured CCFs for mCherry- and mCherry2-mEGFP, and observed a ~40% higher SNR of mCherry2-mEGFP CCFs (SNR_mCh-mEGFP_=1.39±0.10, SNR_mCh2-mEGFP_=1.93±0.10; Figure 5D).

These results demonstrate that using mCherry2 instead of mCherry in cross-correlation experiments leads to a more accurate quantification of the spectral cross-correlation, i.e. of the degree of binding in hetero-interactions or the mobility of hetero-complexes.

## DISCUSSION

In the last decade, FFS-based techniques have become widely used approaches to measure protein dynamics, interactions and oligomerization directly in living cells and organisms^2,6,28,41–44^. One of the most important quantities in these studies is the molecular brightness, i.e. the photon count rate per molecule, which is used as a measure of oligomerization of fluorescently labeled proteins^4^.

In this work, we present a comprehensive analysis of FPs and their suitability for brightness-based oligomerization and cross-correlation interaction studies. Differently from previous reports^4,5^, we consistently obtained lower than expected values for the normalized brightness of homo-dimers for the most common FPs (i.e. mEGFP, mEYFP and mCherry, see Fig. 1A, B). We therefore performed a systematic comparison of commonly used FPs, including also several novel monomeric red FPs (i.e. mCherry2, mRuby3, mCardinal, mScarlet and mScarlet-I), under various conditions. To rule out systematic errors related to the experimental setup or FFS technique used, we performed a combination of pFCS, sFCS and N&B approaches on independent microscopy setups, obtaining reproducible results. Moreover, we excluded potential artifacts deriving from the specific expression system, by comparing different cellular compartments (cytoplasm, nucleus, PM), cell types (HEK 293T, A549, CHO-K1, HeLa) and temperatures (23°C, 37°C), as shown in Figs. 1 and S1. By performing FLIM measurements of mEGFP (Fig. S2), we ruled out the presence of multiple brightness states that might lower homo-dimer brightness values^17^, or energy transfer to non-fluorescent states of mEGFP dimers, in agreement with previous studies^13^. We thus conclude that the observed brightness decrease of FP dimers indicates the presence of a non-fluorescent protein fraction, independent of the experimental conditions. This conclusion is supported by previous reports discussing FP specific photophysical transitions (e.g. blinking, flickering, long-lived dark states)^12–15,17,18,45^ and maturation times^30^. For EGFP, dark state fractions of 20-40% were reported *in vitro*, depending on pH and excitation power^13^. These values agree well with the 30-35% of non-fluorescent mEGFPs that we observed directly in living cells.

We next investigated how the presence of non-fluorescent states exactly affects brightness data of protein complexes of known oligomeric state. This information can then be used to correctly determine the oligomerization state of an unknown protein in general. To this aim, we measured the brightness of mEGFP-homo-oligomers as well as two Influenza A virus HA protein variants with different oligomeric states: HA-wt and an HA-TMD mutant. Biochemical studies have shown that the latter proteins assemble as trimers and dimers, respectively^33–35^. We observed a systematic underestimation of the brightness for all samples compared to the expected values (Figs. 2, 3) and showed that a simple two-state model, determining the p_f_ for each FP from the homo-dimer brightness, successfully yields correct estimates of the oligomeric state. Since the assumption of a constant p_f_ obtained for 2xmEGFP reproduces the correct oligomeric state of higher oligomers, we conclude that maturation is constant for each single FP-subunit within a certain oligomer. In other words, it is sufficient to know the brightness of a FP monomer and homo-dimer, in order to quantify the oligomeric state of larger complexes. It is worth emphasizing that this procedure works well not only for mEGFP-homo-oligomers, but also for large self-assembling protein complexes such as the 12-meric *E.Coli* GlnA^32^, and transmembrane proteins such as the Influenza A virus HA. An equivalent correction approach was used before in single molecule subunit counting studies, albeit mostly restricted to (m)EGFP^10,11,21^. Our results clearly show that a precise correction of the non-fluorescent FP fraction and knowledge of the p_f_ for all involved FPs are absolutely necessary for a correct quantification of protein oligomerization in FFS techniques. Ignoring non-fluorescent FPs leads to a strong misinterpretation of the data, e.g. a tetramer being classified as a trimer (Fig 2). These systematic errors are particularly pronounced for FPs with a low p_f_, as found e.g. for mCherry (~40%), a FP often used in the past to determine the stoichiometry of protein complexes^2,46^. Moreover, FPs possessing low p_f_ severely suffer from low dynamic ranges, since the brightness increase per FP-subunit is only marginal (see Figs. 1, 4), e.g. a mCherry tetramer would be only 2.2 times brighter than a monomer. Nevertheless, contradictory results are reported in this context by studies employing FFS techniques. While very few studies confirm the presence of non-fluorescent mEGFP fractions^22^, others report dimer brightness values of 2 (i.e. the absence of a non-fluorescent FP fraction)^4,5^. However, the latter studies were all performed with two-photon excitation, which may influence the transition to non-fluorescent states^17^. In this context, our data provide the first complete and systematic comparison of p_f_ for several FPs, in one-photon excitation setups.

In order to measure multiple species simultaneously, red FPs are required due to their spectral separation from mEGFP. Additionally, they are a preferential choice for tissue and animal imaging, due reduced light absorption and autofluorescence in the red and far-red spectral region^29^. Given the suboptimal p_f_ we determined for mCherry, we screened several recently developed monomeric red FPs^36–39^. The p_f_ is in fact an essential parameter that, until now, has not received appropriate attention in reports of new FPs. The suitability of these proteins for FFS studies depends on three important fluorophore characteristics: 1) a high photostability is required to enable temporal measurements under continuous illumination, 2) a high molecular brightness is needed to obtain a signal-to-noise ratio sufficient to detect single-molecule fluctuations, 3) a high p_f_ is essential for a maximal dynamic range that allows reliable oligomerization measurements. Thus, red FPs which fulfill only one or two of these requirements are not recommended for FFS measurements. Among all red FPs investigated in this study, we found only one fulfilling all three important criteria: mCherry2, a rarely used mCherry variant^36^ that has not been entirely characterized yet. However, for the remaining red FPs tested here, we found either low photostability albeit high monomer brightness (mRuby3, mScarlet, mScarlet-I; Fig. 4A, B), and/or low to medium p_f_ of 20-45% (mCardinal, mCherry, mRuby3, mScarlet; Fig. 4C), very similar to previously published values for mRFP^24,25^ and mCherry^25^. In contrast, mCherry2 possesses a high p_f_ of ~70%. Very recent studies of FP maturation times report a faster maturation of mCherry2 and mScarlet-I compared to mCherry/mScarlet^30^. Together with our findings, this indicates that faster FP maturation could be the reason for the observed higher p_f_.

Finally, we demonstrate that quantification of hetero-interactions via cross-correlation approaches, so far typically performed with mCherry^1,2,40^, can be substantially improved by using mCherry2 instead. In agreement with the reported similar p_f_ of mEGFP and mCherry2 (Figs. 1, 4), we observed that the amount of hetero-dimers containing both fluorescent mEGFP and mCherry2 increased significantly compared to those containing mCherry. For this reason, the CCF signal-to-noise ratio for mCherry2-mEGFP complexes increased by 40% compared to that measured for mCherry-mEGFP hetero-dimers (Fig. 5D). This could be particularly relevant for investigations of weak interactions, in which only a small number of hetero-complexes is present, compared to the vast amount of non-interacting molecules. Additionally, crosscorrelation techniques have been recently applied in living multicellular organisms^6,28^, which require low illumination to avoid phototoxicity and thus generally suffer from low signal-to-noise ratios. Therefore, we recommend using mCherry2 as the novel standard red FP in brightness and cross-correlation measurements.

In conclusion, this study provides a useful, comprehensive resource for applying FFS techniques to quantify protein oligomerization and interactions. We provide a clear, simple methodology to test and correct for the presence of non-fluorescent states, and argue that such controls should become a prerequisite in brightness-based FFS studies to avoid systematic errors in the quantification of protein oligomerization. Finally, our results suggest that the apparent fluorescence probability is an important fluorophore characteristic that should be considered and reported when developing new FPs and we provide a simple assay to determine this quantity.

## ACKNOWLEDGEMENTS

This work was supported by the Deutsche Forschungsgemeinschaft (DFG) grants (254850309 to S.C. and SFB 740, TP C3 to A.H.).

## MATERIALS AND METHODS

### Cell culture

Human embryonic kidney (HEK) cells from the 293T line (purchased from ATCC^®^, CRL-3216™), human epithelial lung cells A549 (ATCC^®^, CCL-185™), chinese hamster ovary (CHO) cells from the K1 line (ATCC^®^, CCL-61™), human epithelial cervix cells HeLa (ATCC^®^, CCL-2™) and human bone osteosarcoma epithelial cells U2OS (a kind gift from Ana García Sáez, University of Tübingen) were cultured in Dulbecco’s modified Eagle medium (DMEM) with the addition of fetal bovine serum (10%) and L-Glutamine (4 mM). Cells were passaged every 3-5 days, no more than 15 times. All solutions, buffers and media used for cell culture were purchased from PAN-Biotech (Aidenbach, Germany).

### Fluorescent protein constructs

For the cloning of all following constructs, standard PCRs with custom-designed primers were performed to obtain monomeric FP cassettes, followed by digestion with fast digest restriction enzymes and ligation with T4-DNA-Ligase according to the manufacturer’s instructions. All enzymes were purchased from Thermo Fisher Scientific, unless specified otherwise.

The constructs 2xmEGFP, 3xmEGFP and 4xmEGFP (i.e. mEGFP dimer, trimer and tetramer) were obtained by step-wise cloning of monomeric mEGFP cassettes amplified from mEGFP-N1, a gift from Michael Davidson (Addgene plasmid #54767). First, 2xmEGFP was generated by ligating an mEGFP cassette into mEGFP-N1 digested with BamHI and AgeI. Subsequently, an additional monomeric mEGFP cassette was ligated into 2xmEGFP by digestion with KpnI and BamHI to generate 3xmEGFP. Finally, 4xmEGFP was obtained by ligation of an additional monomeric mEGFP cassette into 3xmEGFP by digestion with EcoRI and KpnI. All mEGFP subunits are linked by a polypeptide sequence of five amino acids. To ensure purity of mEGFP-homo-oligomers, all full-length inserts were subcloned into pcDNA™3.1(+) (Thermo Fisher Scientific) possessing ampicillin instead of kanamycin resistance.

The GlnA-mEGFP plasmid was a kind gift from Ana García Sáez (University of Tübingen) and cloned based on pGlnA-Ypet (gift from Mike Heilemann, Addgene plasmid #98278).

To obtain 2xmEYFP, mEYFP was amplified from mEYFP-N1^1^ and inserted into mEYFP-C1 by digestion with KpnI and BamHI.

The plasmids mCherry-C1 and mCherry2-C1/N1 (gifts from Michael Davidson, Addgene plasmids #54563 and #54517, respectively) were used to generate 2xmCherry and 2xmCherry2, respectively. First, mCherry-C1 and –N1 were generated by amplification of mCherry from mCherry-pLEXY plasmid (a gift from Barbara Di Ventura & Roland Eils, Addgene plasmid #72656) and inserted into a pBR322 empty vector. A second mCherry cassette was inserted into this vector by digestion with XhoI and BamHI to obtain 2xmCherry (i.e. mCherry dimer). The 2xmCherry2 (mCherry2 dimer) plasmid was generated by amplification of mCherry2 and insertion of this construct into mCherry2-C1 through digestion with XhoI and BamHI. To clone mRuby3-C1, mRuby3 was amplified from the pKanCMV-mClover3-mRuby3 plasmid, a gift from Michael Lin (Addgene plasmid #74252). The obtained PCR product was digested with AgeI and XhoI and exchanged with mEYFP from digested mEYFP-C1 plasmid. For 2xmRuby3 (mRuby3 dimer), mRuby3 was again amplified by PCR and the product inserted into mRuby3-C1 by digestion with KpnI and BamHI. The mCardinal-C1/N1 plasmids were a gift from Michael Davidson (Addgene plasmids #54590 and #54799). To obtain 2xmCardinal (mCardinal dimer), mCardinal was amplified from mCardinal-C1 and the PCR product inserted into mCardinal-C1 by digestion with KpnI and BamHI. The plasmids mScarlet-C1 and mScarlet-I-C1 are gifts from Dorus Gardella (Addgene plasmids #85042 and #85044). 2xmScarlet (mScarlet dimer) and 2xmScarlet-I (mScarlet-I dimer) were generated by amplification of mScarlet and mScarlet-I from the corresponding plasmids and reintegration into mScarlet-C1 and mScarlet-I-C1 by digestion with XhoI and KpnI. To ensure purity of dimers, all full-length dimers were subcloned into pcDNA™3.1(+) (Thermo Fisher Scientific) possessing ampicillin instead of kanamycin resistance.

The hetero-dimers mCherry-mEGFP and mCherry2-mEGFP were generated by amplification of mCherry and mCherry2, respectively, and insertion of the obtained constructs into mEGFP-C1, (Michael Davidson, Addgene plasmid #54759), by digestion with XhoI and BamHI. Both fluorophores are linked by five and seven amino acids, respectively.

The membrane constructs consisting of mEGFP linked to a myristoylated and palmitoylated peptide (mp 1xmEGFP) and its dimer mp 2xmEGFP were kind gifts from Richard J. Ward (University of Glasgow)^2^. The analogue mp 1xmEYFP construct was obtained as described elsewhere^1^. To generate mp 2xmEYFP, the 2xmEYFP cassette described above was transferred into a myr-palm-mCardinal vector^3^, by digestion with AgeI and BamHI. The GPI mCherry (glycosylphosphatidylinositol-anchored mCherry) plasmid was a kind gift from Roland Schwarzer (Gladstone Institute, San Francisco). Based on this plasmid, GPI 2xmCherry was generated by amplification of a mCherry cassette and ligation of the obtained insert into GPI mCherry digested, using SalI and BamHI.

The Influenza virus A/chicken/FPV/Rostock/1934 hemagglutinin (HA) constructs HA-wt-mEGFP and HA-TMD-mEGFP were cloned based on the previously described HA-wt-mEYFP^1^ and HA-TMD-mEYFP^4^ plasmids. HA-wt-mEYFP contains full-length HA protein fused to mEYFP at the C-terminus, whereas in HA-TMD-mEYFP a large part of the extracellular domain of HA is replaced by mEYFP. To clone HA-wt-mEGFP, HA-wt-mEYFP was digested using BglII and SacII (New England Biolabs) and the obtained HA insert ligated into mEGFP-N1. For HA-TMD-mEGFP, HA-TMD-mEYFP plasmid and mEGFP-N1 vector were digested with AgeI and BsrGI to replace mEYFP with mEGFP.

All plasmids generated in this work will be made available on Addgene.

### Preparation for Microscopy Experiments

For microscopy experiments, 6×10^5^ (HEK) or 4×10^5^ (A549, CHO, HeLa) cells were seeded in 35 mm dishes (CellVis, Mountain View, CA or MatTek Corp., Ashland, MA) with optical glass bottom, 24 h before transfection. HEK 293T cells were preferred for scanning FCS (sFCS) measurements since they are sufficiently thick and therefore ideal for sFCS based data acquisition perpendicular to the PM. A549 cells are rather flat and characterized by a large cytoplasmic volume that is more suitable for point FCS measurements in the cytoplasm. Cells were transfected 16-24 h prior to the experiment using between 200 ng and 1 μg plasmid per dish with Turbofect (HEK, HeLa, CHO) or Lipofectamin3000 (A549) according to the manufacturer’s instructions (Thermo Fisher Scientific). Briefly, plasmids were incubated for 20 min with 3 μl Turbofect diluted in 50 μl serum-free medium, or 15 min with 4 μl P3000 per 1 μg plasmid and 2 μl Lipofectamine3000 diluted in 100 μl serum-free medium, and then added dropwise to the cells.

### Confocal Microscopy System

Confocal imaging and point Fluorescence (Cross-) Correlation Spectroscopy (pF(C)CS) measurements were performed on an Olympus FluoView FV-1000 system (Olympus, Tokyo, Japan) using a 60x, 1.2NA water immersion objective. Scanning Fluorescence (Cross-) Correlation Spectroscopy (sF(C)CS) and Number&Brightness (N&B) measurements were performed on a Zeiss LSM780 system (Carl Zeiss, Oberkochen, Germany) using a 40x, 1.2NA water immersion objective. Samples were excited with a 488 nm Argon laser (mEGFP, mEYFP) and a 561 nm (Zeiss instrument) or 559 nm (Olympus) diode laser (mCherry, mCherry2, mCardinal, mRuby3, mScarlet, mScarlet-I). For measurements with 488 nm excitation, fluorescence was detected between 500 and 600 nm, after passing through a 488 nm dichroic mirror, using SPAD (PicoQuant, Berlin, Germany, mounted on Olympus instrument) or GaAsP (Zeiss instrument) detectors. For 561 nm or 559 nm excitation, fluorescence emission passed through a 488/561 nm (Zeiss) or 405/488/559/635 nm (Olympus) dichroic mirror and was detected between 570 and 695 nm (Zeiss) or using a 635 nm long-pass filter (Olympus). For pFCCS measurements, fluorophores were excited using 488 nm and 559 nm laser lines. Excitation and detection light were separated using a 405/488/559/635 nm dichroic mirror. Fluorescence was separated on two SPAD detectors using a 570 nm dichroic mirror and detected after passing through a 520/35 nm bandpass filter (mEGFP channel) or a 635 nm long-pass filter (mCherry or mCherry2 channel) to minimize cross-talk.

### Fluorescence (Cross-) Correlation Spectroscopy

Point F(C)CS measurements were routinely performed for 90 s and recorded using the SymPhoTime64 software (PicoQuant GmbH, Berlin, Germany). Laser powers were adjusted to keep photobleaching below 20%. Typical values were ~3.3 μW (488 nm) and ~6 μW (559 nm). The size of the confocal pinhole was set to 90 μm. PicoQuant ptu-files containing recorded photon arrival times were converted to intensity time series and subsequently analyzed using a custom-written MATLAB Code (The MathWorks, Natick, MA, USA). First, the intensity time series was binned in 5 μs intervals. To correct for signal decrease due to photobleaching, the fluorescence time series was fitted with a two-component exponential function, and a correction was applied^5^. Then, autocorrelation functions (ACFs) and, in case of two-color experiments (g=green channel, r=red channel), cross-correlation functions (CCFs) were calculated as follows, using a multiple tau algorithm:

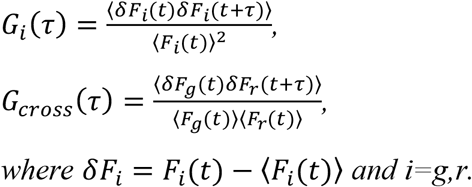

To avoid artefacts caused by long-term instabilities or single bright events, CFs were calculated segment-wise (10 segments) and then averaged. Segments showing clear distortions were manually removed from the analysis.

A model for anomalous three-dimensional diffusion and a Gaussian confocal volume geometry was fitted to the ACFs^6^:

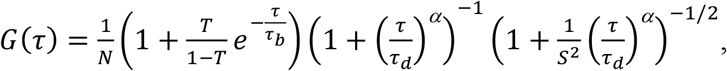

where the exponential term accounts for photophysical transitions of a fraction T of fluorescent proteins. The parameter τ_b_ was constrained to values lower than 50 μs for mEGFP^7^ or mEYFP^8^ and 200 μs for mCherry/mCherry2^9^. The anomaly parameter a was introduced to account for anomalous subdiffusion of proteins in the cytoplasm^6^ and constrained to values between 0.5 and 1. The particle number N and diffusion time τ_d_ were obtained from the fit. To calibrate the focal volume, pFCS measurements with Alexa Fluor® 488 or Rhodamine B dissolved in water at 50 nM were performed at the same laser power. The structure parameter S was fixed to the value determined in the calibration measurement (typically around 4 to 8). The molecular brightness was calculated by dividing the mean count rate by the particle number determined from the fit.

For two-color measurements, all ACFs were used to fit the diffusion model described above. Relative cross-correlation values were calculated from the amplitudes of ACFs and CCFs: 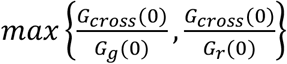, where *G_cross_*(0) is the amplitude of the CCF and *G_i_*(0) is the amplitude of the ACF in the *i*-th channel. The signal-to-noise ratio (SNR) of the CCFs was calculated by summing the cross-correlation values divided by their variance over all points of the CCF. The variance of each point of the CCF was calculated by the multiple tau algorithm^10^.

### Scanning Fluorescence Correlation Spectroscopy

For one-color sFCS measurements, a line scan of 128×1 pixels (pixel size 160 nm) was performed perpendicular to the membrane with 472.73 μs scan time. Typically, 250,000-500,000 lines were acquired (total scan time 2 to 4 min) in photon counting mode. Laser powers were adjusted to keep photobleaching below 20%. Typical values were ~1.8 μW (488 nm) and ~6 μW (561 nm). Scanning data were exported as TIFF files, imported and analyzed in MATLAB (The MathWorks, Natick, MA) using custom-written code. sFCS analysis follows the procedure described previously^3,11^. Briefly, all lines were aligned as kymographs and divided in blocks of 1000 lines. In each block, lines were summed up column-wise and the x position with maximum fluorescence was determined. This position defines the membrane position in each block and is used to align all lines to a common origin. Then, all aligned line scans were averaged over time and fitted with a Gaussian function. The pixels corresponding to the membrane were defined as pixels which are within ±2.5σ of the peak. In each line, these pixels were integrated, providing the membrane fluorescence time series F(t). When needed, a background correction was applied by subtracting the average pixel fluorescence value on the inner side of the membrane multiplied by 2.5σ (in pixel units) from the membrane fluorescence, in blocks of 1000 lines^12^. In order to correct for depletion due to photobleaching, the fluorescence time series was fitted with a two-component exponential function and a correction was applied^5^. Finally, the ACF was calculated as described above.

A model for two-dimensional diffusion in the membrane and a Gaussian focal volume geometry^11^ was fitted to the ACF:

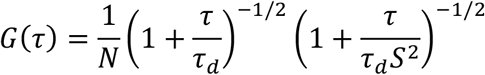

The focal volume calibration was performed as described for pF(C)CS. Diffusion coefficients (D) were calculated using the calibrated waist of the focal volume, *D* = *ω*_0_^2^/4*τ_d_*. The molecular brightness was calculated by dividing the mean count rate by the particle number determined from the fit: 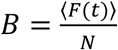.

### Number&Brightness Analysis

N&B experiments were performed as previously described^13^, with a modified acquisition mode. Briefly, 200 images of 128×64-128 pixels were acquired per measurement, using a 300 nm pixel size and 25 μs pixel dwell time. Laser powers were maintained low enough to keep bleaching below 10% of the initial fluorescence signal (typically ~0.7 μW for 488 nm and ~4.9 μW for 561 nm) except for mRuby3 and mScarlet/ mScarlet-I. CZI image output files were imported in MATLAB using the Bioformats package^14^ and analyzed using a custom-written script. Before further analysis, pixels corresponding to cell cytoplasm or nucleus were selected manually as region of interest. Brightness values were calculated as described^13^, applying a boxcar algorithm to filter extraneous long-lived fluctuations^15,16^. Pixels with count rates above 2 MHz were excluded from the analysis to avoid pile-up effects. To further calibrate the detector response, we measured the brightness on a reflective metal surface and dried dye solutions. The thus obtained brightness-versus-intensity plots (which should be constant and equal to 0 for all intensity values^13^) were used to correct the actual experimental data.

### Fluorescence Lifetime Imaging Microscopy

FLIM measurements were performed on an Olympus FluoView FV-1000 system (Olympus, Tokyo, Japan) equipped with a time-resolved LSM upgrade kit (PicoQuant GmbH, Berlin, Germany) using a 60x, 1.2NA water immersion objective. Images of 512×512 pixels per frame were acquired after excitation with a pulsed-laser diode at 488 nm. Fluorescence was detected using a SPAD detector and a 520/35 nm bandpass filter. In each measurement, a minimum of 10^5^ photons were recorded by accumulation of 60 frames over a time period of 90 s. Regions of interest in the cytoplasm of cells were analyzed using SymPhoTime64 software (PicoQuant GmbH, Berlin, Germany) taking into account the instrument response function determined by measuring a saturated Erythrosine B solution according to manufacturer’s instructions. Resulting decay curves were fitted using a mono-exponential function.

### Brightness calibration and fluorophore maturation

The molecular brightness, i.e. the photon count rate per molecule, serves as a measure for the oligomeric state of protein complexes. This quantity is affected by the presence of non-fluorescent FP fractions, which can result from several processes: 1) Photophysical processes such as long-lived dark states, blinking or flickering between an *on* and *off* state, 2) FP maturation, i.e. FPs that have not maturated yet, 3) Incorrectly folded FPs. To quantify the amount of non-fluorescent FPs, we consider all these processes together in a single parameter, the apparent fluorescence probability (pf), i.e. the probability of a FP to emit a fluorescence signal. The fluorescence emitted by an oligomer can then be modeled with a binomial distribution, assuming that each fluorophore monomer emits photons with brightness ε and with a probability p_f_. The probability of detecting a brightness value iε for an n-mer is thus 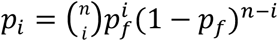. Hence, the ensemble-averaged brightness detected from a number of N n-mers is:

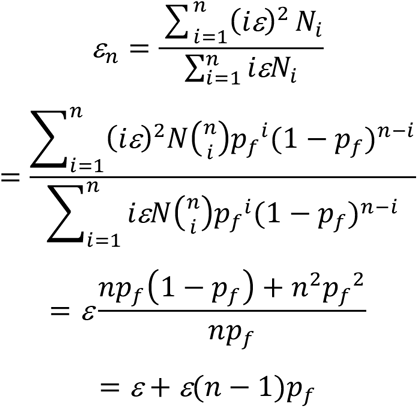

In the analysis, we normalized all brightness values to the median brightness of the corresponding monomer sample measured under the same conditions: 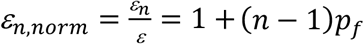. We used the median of the normalized dimer brightness to determine the probability p_f_ for each construct, *p_f_* = *ε_2,norm_* − 1. We can now invert the equation for the n-mer brightness to calculate the true oligomeric state, i.e. the brightness if all subunits were constantly fluorescent, from the measured brightness ε_n_: 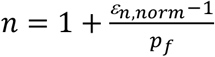.

We applied this transformation to every brightness data point and obtained the “corrected” brightness. Notably, this transformation holds true also for fluorophores which have two brightness states rather than an *on* and *off* state^17^.

### Statistical analysis

All data are displayed as box plots indicating the median values and whiskers ranging from minimum to maximum values. Statistical significance was tested using a two-tailed Mann-Whitney test or one-way ANOVA for multiple comparisons. Quantities in the main text are given as mean±S.E.M.

### Code availability

MATLAB custom-written code is available upon request from the corresponding author.

### Data availability

The datasets analyzed during the current study are available from the corresponding author on reasonable request.

